# Using Google Trends to Examine the Spatio-Temporal Incidence and Behavioral Patterns of Dengue Disease: A Case Study in Metropolitan Manila, Philippines

**DOI:** 10.1101/424630

**Authors:** Howell T. Ho, Thaddeus M. Carvajal, John Robert Bautista, Jayson Dale R. Capistrano, Katherine M. Viacrusis, Lara Fides T. Hernandez, Kozo Watanabe

## Abstract

Dengue is a major public health concern and an economic burden in the Philippines. Despite the country’s improved dengue surveillance, it still suffers from various setbacks and therefore needs to be complemented with alternative approaches. Previous studies have demonstrated the potential of internet-based surveillance such as Google Dengue Trends (GDT) in supplementing current epidemiological methods for predicting future dengue outbreaks and patterns. With this, our study aims to assess the temporal relationship of GDT and dengue incidence in Metropolitan Manila from previous years and examine web search behavior of the population towards the disease. The study collated and organized the population statistics and reported dengue cases in Metropolitan Manila from respective government agencies to calculate the spatial and temporal dengue incidence. The relative search volume of the term ‘dengue’ and top dengue-related search queries in Metropolitan Manila were obtained and organized from the Google trends platform. Data processing of GDT and dengue incidence was performed by conducting an ‘adjustment’ procedure and subsequently used for correlation and cross-correlation analyses. Moreover, a thematic analysis was employed on the top dengue-related search queries. Results revealed a high temporal relationship between GDT and dengue incidence when either one of the variables is adjusted. Cross-correlation showed that there is delayed effect (1-2 weeks) of GDT to dengue incidence, demonstrating its potential in predicting future dengue outbreaks and patterns in Metropolitan Manila. Thematic analysis of dengue-related search queries indicated 5 categories namely; (a) dengue, (b) sign and symptoms of dengue, (c) treatment and prevention, (d) mosquito and (e) other diseases where the majority of the search queries was ‘signs and symptoms’ which indicate the health-seeking behavior of the population towards the disease.

## 1. Introduction

Dengue is one of the leading and most important mosquito-borne viral diseases in the world where 2.5 billion people worldwide is estimated to be at risk of contracting the disease [1]. Southeast Asia has been identified to be highly vulnerable to mosquito-borne diseases[2] where incessant massive outbreaks of dengue have been reported in selected countries [3,4,5,6,7,8]. Therefore, the significant health and economic burden [9] brought by dengue in the Philippines make it a major public health concern leading towards becoming a national notifiable disease since 1958 [10].

The National Epidemiology Centre (NEC) of the Department of Health in the Philippines has developed a reporting system for dengue in all its disease reporting units, such as hospitals and rural health facilities [11]. For the past decade, this traditional, healthcare-based, and government-implemented surveillance system has undergone improvements, yet it still suffers from untimely reporting, aggregation, and publishing of data. Due to surveillance problems, the identification and efficiency of intervention that may prevent dengue epidemics are grossly limited. Many suggestions involving alternative approaches outside the virological and clinical domains (e.g. telephone triage calls, sales of over-the-counter drugs, and school/work absenteeism) have been made in parallel with traditional surveillance [12,13,14], Among these non-traditional suggestions, online activity or internet search tracking has shown the potential to complement current epidemiological methods because of its consistency, efficiency, and manifestation of real-time population trends [15,16,17,18,19].

Internet access and use have increased globally, including the Philippines. The internet, together with social media, has become capable of facilitating disease surveillance, mass communication, health education, and knowledge translation and collaboration amongst health providers [20]. Google Trends is an accessible internet platform that provides geospatial and temporal patterns of search volumes for user-specified terms towards public health surveillance. The association and predictive power of Google Trends towards disease surveillance [21] have been studied extensively especially for diseases such as flu [22], HIV [23], scarlet fever [24], malaria [25], ebola [26], and Zika [27]. Countries like Bolivia, Brazil, India, Indonesia, Singapore, Mexico, and Valenzuela have recently investigated the application of Google Trends for dengue surveillance and it has been observed to be highly correlated with the temporal pattern of dengue in a country-wide scale [16,17,28,29]. Therefore, utilizing this platform may present a valuable complement in assisting real-time dengue case surveillance [16] and can be potentially extended towards predicting early dengue disease outbreaks [17]. While majority of the previous studies investigated the monthly association of Google Dengue trends (GDT) and reported dengue cases, it left us an impression that present GDT values can be adapted immediately for associative or predictive purposes. However, this is not the case as GDT can only provide weekly or daily values with a limited time period. For instance, weekly values can only be obtained for a maximum range of 4 years [30]. Beyond that, the resulting GDT values will expand in monthly intervals. ‘Stitching’ yearly GDT weekly values is not recommended since these are relative and are based on the highest query volume for a given time period. If researchers were not keen on such limitation, reporting the association or predictive power of GDT may prove to be an under-estimated value. For this reason, there is a need to adjust and normalize GDT values in order to capture and provide the correct estimation of its associative temporal pattern [31].

Another potential advantage of Google Trends is the understanding of health-seeking behaviors of the targeted population toward specific diseases [20]. It can present significant insights into the population behavior and health-related phenomena of diseases [32]. Because of its capacity to reflect related important topics from search queries at a given time period, Google Trends provides valuable data on quantifying the health-seeking behavior of a specified population [33]. As to date, there have been no studies that evaluated related-search queries of dengue. Therefore, investigating dengue-related queries may provide a better understanding of the population’s health-seeking behaviors regarding the disease [34].

The primary aim of the study is to utilize Google Trends search volume data (GDT) to examine Dengue patterns in Metropolitan Manila, Philippines. Specifically, it aims to accomplish two objectives: (1) to assess the temporal relationship between Google Dengue Trends (GDT) and dengue incidence, and (2) to examine the web search behavior of users towards dengue.

## 2. Materials and Methods

### 2.1. Study Area and Population Demographics

Metropolitan Manila is also known as the National Capital Region (NCR) of the Philippines. It is located at the Eastern shore of Manila Bay in Southwestern Luzon, Philippines (14°50’ N, 121 °E) with an area of 636 km^2^. The region contains 16 cities and one municipality with a total population of 12.8 million [35] (Figure S1). The map layer of the administrative city boundaries of Metropolitan Manila was obtained from the Philippine GIS Data clearinghouse [36]. The population statistics of each city and municipality were obtained from the Philippine Statistics Authority agency [35]. The Philippine population census was only reported during the years 2010 and 2015. Thus, we utilized the compounded population growth rate to calculate the population of the entire region and each city/municipality for the years 2009, 2011, 2012, 2013 and 2014 with the assumption of a fixed growth rate.

### 2.2. Data Source and Processing

Reported dengue cases of Metropolitan Manila (1st Morbidity week of 2009 until 52nd week of 2014) were obtained from the National Epidemiology Center of the Department of Health. Incidence rate of dengue was calculated by dividing the number of cases to the total population size for a given year multiplied by a factor of 1,000. Dengue incidence (DI) was computed (a) annually per city (Figure 2a-f) and (b) weekly in the entire region. Subsequently, we adjusted the weekly values of dengue incidence (AdjDI) by dividing the observed value to the maximum observed value of the year. Furthermore, log transformation was also performed on both (e.g. observed, adjusted) values in order to reduce the skewing of the data and destabilize the variance.

Search query data from Google Trends can be mined from their website (https://trends.google.com/trends) using the term “dengue”. For the purposes of this study, we abbreviate the Google Dengue Trends in the Philippines as “GDT”. This platform shows the relative search volume (RSV) which is the query share of a term (e.g. dengue) for a given time period and location. The value ranges from 0 -100 which are normalized from the highest query share of the term over the time series. This value also represents the “search interest” index where a term value of 100 is denoted as the peak popularity while a value of 50 represents exactly half of the popularity. On the other hand, a term with a value of zero means there are insufficient data. GDT limits the reported weekly values for a certain period of time and in order to resolve this issue, we adapted the adjusting procedure of Risteski and Davcev’s [30,31]. First we obtained the monthly-level GDT query of dengue for the entire period of 2009-2014 and, then collected GDT weekly-level query per year from 2009-2014. Afterwards, we multiplied each weekly-level value by the monthly-level query result and divided it to the monthly-level average from the weekly data. This adjustment procedure accounts for differences in the relative prevalence of searches over time in the stacked weekly-level data.

Additional data displayed are the cities during the previously mentioned time period in Metropolitan Manila that search for the term ‘dengue’. GDT also displays words and phrases referred to as “related queries” of dengue. These queries are divided into (a) top and (b) rising categories. The top category refers to the most popular search queries and contains score values (0-100). While, the rising category refers to queries with the biggest increase in search frequency since the last time period. A select number of search queries can be labelled as “breakout” referring to a tremendous increase at that given time.

### 2.3. Analysis

First, Pearson correlation was performed to correlate data types of dengue and GDT. Moreover, cross-correlation analysis was performed between the two time series data from 0–25 week lags. In Computations were done using the stats package of the R program version 3.3.5 [37]. Second, we descriptively analyzed the GDT values in all cities over each time period with respect to the spatial dengue incidence of Metropolitan Manila. Heat maps were generated using ArcGIS version 10.2 [38].

The top and rising dengue-related search queries for each period were collected and organized. Thematic analysis was performed on the search queries listed in the top query category. To quantify the categorized themes per year, the percentage value was used by obtaining the sum of GDT values for each year and category and then dividing it to the maximum GDT summation value of a particular year. Word clouds of the top search queries in each period were created using Wordart (www.wordart.com). Previous studies have used this web application to create customized word clouds wherein the size of the words or phrases can be adjusted based on their frequency [39,40]. However, instead of frequency, the size of each search query was based on the ‘interest over time’ value provided by Google Trends. Simply put, bigger words in the word cloud will mean high search query value and in turn, will mean high interest over time. In this study, the word cloud was patterned using the map of Metropolitan Manila.

## 3. Results

### 3.1. Association of GDT and Dengue incidence

Figure 1 shows the temporal pattern of the observed and adjusted values of dengue and GDT. It can be observed that the GDT trend (Figure 1a and 1b) follows the seasonal dengue pattern of Metropolitan Manila. However, 2009, 2013, and 2014 GDT trend pattern do not closely match the expected dengue pattern except when it is transformed into its logarithmic function. Further analysis demonstrated that the association of observed weekly values of dengue incidence (DI) were moderately associated (r = 0.405) to the weekly GDT values and the association between GDT and the log transformed DI was slightly lower (r = 0.394) (Table 1).

**Figure 1.**
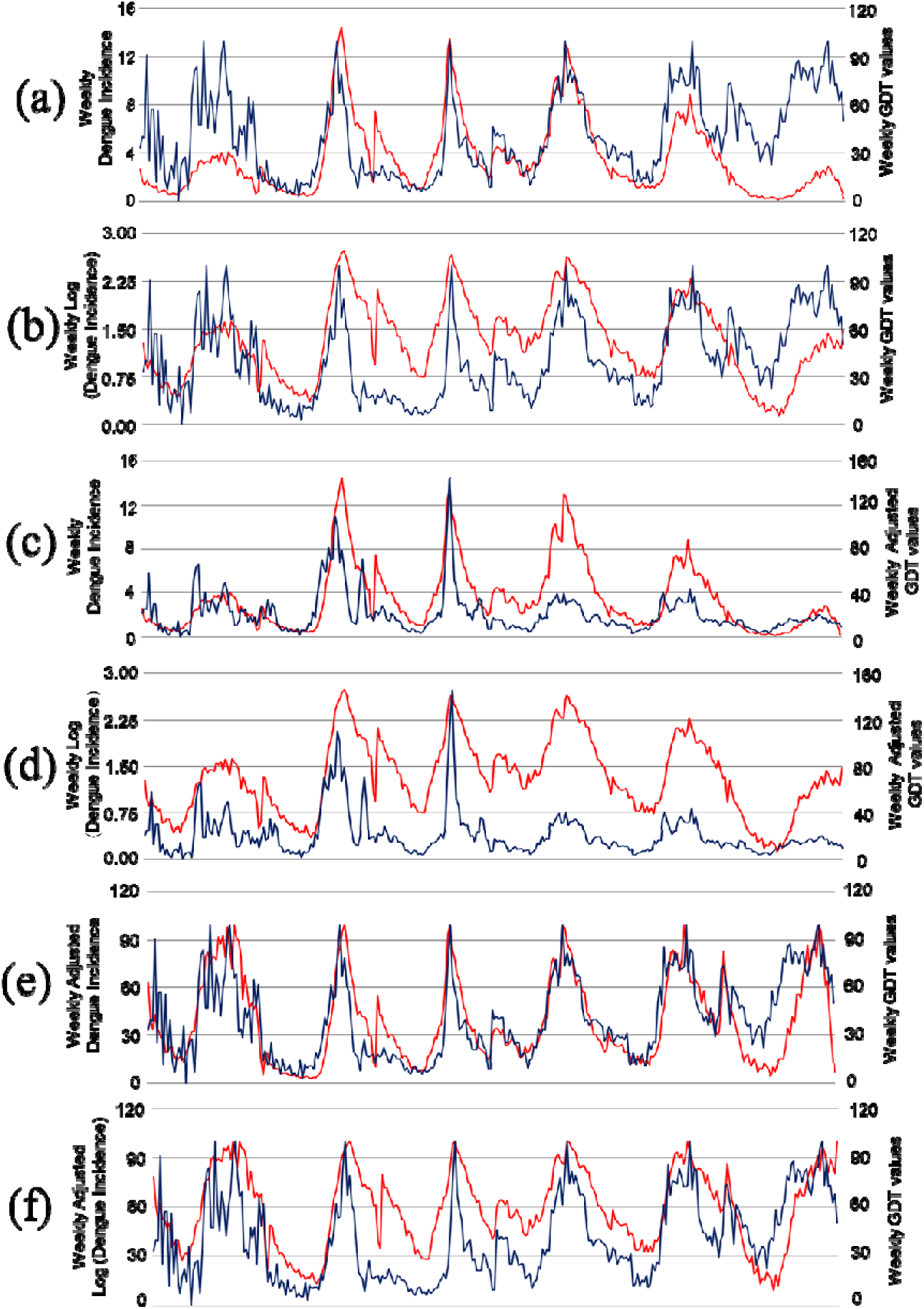
Weekly reports of Dengue (red line) and Google Dengue Trends (blue line) from 2009-2014. (a) Dengue Incidence and GDT values; (b) Log-transformed Dengue Incidence and GDT values; (c) Dengue Incidence and Adjusted GDT values; (d) Log-transformed Dengue Incidence and Adjusted GDT values; (e) Adjusted Dengue Incidence and GDT values; (f) Adjusted Log-transformed Dengue Incidence and GDT values

**Table 1.** Correlation analysis of Dengue and Google Dengue Trends (GDT) and its delayed effects (R^2^)

| Dengue Values | Google Dengue Trend Values |  |  |  |
| --- | --- | --- | --- | --- |
|  | Google Dengue Trend |  | Adjusted Google Dengue Trend |  |
|  | *r* | *lag week ($R^2$)* | *r* | *lag week ($R^2$)* |
| Dengue Incidence | 0.405 | 1 (0.166) | 0.662 | 1 (0.465) |
| Log (Dengue Incidence) | 0.394 | 1 (0.162) | 0.597 | 2 (0.385) |
| Adjusted Dengue Incidence | 0.747 | 1 (0.570) | 0.529 | 2 (0.305) |
| Log (Adjusted Dengue Incidence) | 0.576 | 1 (0.342) | 0.470 | 2 (0.245) |

As per Risteski and Davcev’s [30,31] recommendations, the weekly GDT values (AdjGDT) were adjusted accordingly. It can be observed that the AdjGDT values still follow the seasonal dengue pattern except for the periods of 2012 and 2013 and the trend pattern of the AdjGDT does not conform to the transformed logarithm of dengue incidence (Figure 1c and 1d). The association of the AdjGDT values showed higher correlation coefficients for DI (r= 0.662) and log transformed DI (r= 0.597) as compared to when GDT was not adjusted (Table 1). Interestingly, when the DI was adjusted (AdjDI and logAdjDI) and plotted with the non-adjusted GDT values, the temporal trends were nearly similar to each other (Figure 1e and 1f). Moreover, the association of GDT to AdjDI (r = 0.747) and logAdjDI (r = 0.660) were higher as compared to non-adjusted variables and when AdjGDT was only used (Table 1). Lastly, when both variables were adjusted and plotted, the temporal patterns did not closely match each other (data not shown). This observed patterns are the same when both variables are not adjusted (Figure 1a - 1b). However, correlations of AdjGDT with AdjDI (r = 0.529) and logAdjDI (r = 0.470) were higher as compared to non-adjusted variables.

In addition, Table 1 shows the cross-correlation analysis (lag effects) (Supplemental Material Table S1) of GDT to dengue incidence. Majority of all cross correlation analyses revealed the highest lag association to be at lag 1 and lag 2. This can be visually observed in selected high peaks of both dengue and GDT in Figure 1. For example, in Figure 1e (AdjDI and GDT), prior to the highest dengue incidence peak in 2010, a high GDT peak occurred a week before.

### 3.2. Spatial pattern of GDT and related queries for Dengue

Figure 2 shows the GDT values for Metropolitan Manila cities from 2009 – 2014. It is noteworthy to mention that the cities with GDT increased from two in 2009 to seven in 2013. Makati consistently had such data for the entire 6-year period. This is followed by cities, such as Quezon (5yrs), Manila (5yrs), Pasig (3yrs), Mandaluyong (3yrs), Paranaque (2yrs), and Las Piñas (1yr). Comparing the spatial maps of dengue incidence (Figure 2a-f) and GDT values per city showed no discernable pattern revealing high dengue incidence in a city with high GDT value. Further analysis showed a very low and non-significant relationship (r= 0.223, p=0.283).

**Figure 2.**
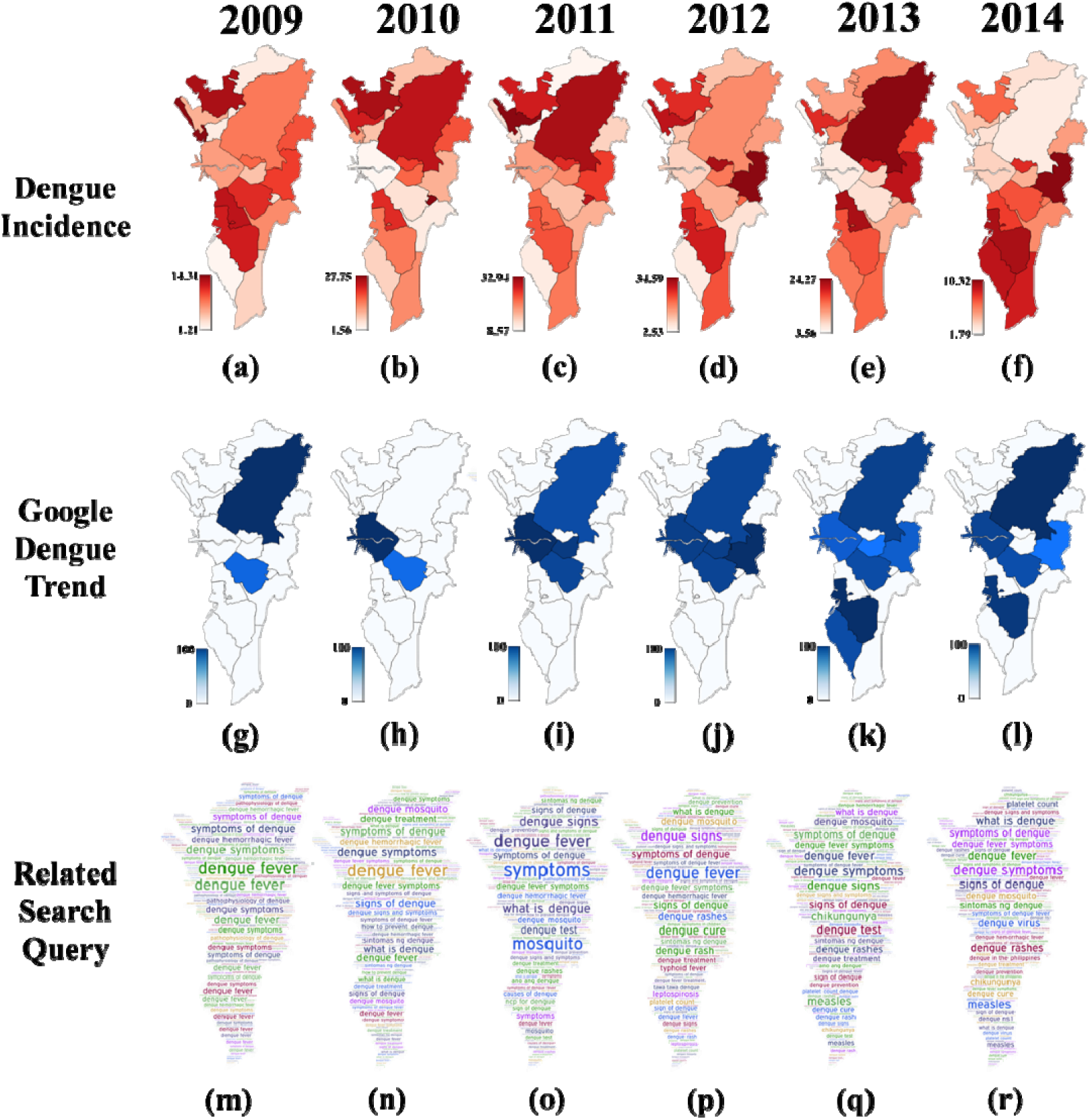
Spatio-temporal pattern of Dengue incidence (a-f), Google Dengue Trends (g-1) and Word Clouds maps of Dengue related search query (m-r)

Table 2 shows that the top search queries from 2009 to 2014 and were categorized under five groups. These were (1) dengue, (2) signs and symptoms, (3) treatment and prevention, (4) mosquito, and (5) other diseases. Based on the GDT percentages across each category and year, many of the listed top queries were related to dengue and its symptoms. Search queries under “dengue” were more popular in 2009. Search queries under this group consisted of phrases related to dengue’s etiology (e.g., causes of dengue and dengue virus) and alternative names (e.g., dengue hemorrhagic fever and dengue fever).

**Table 2.**
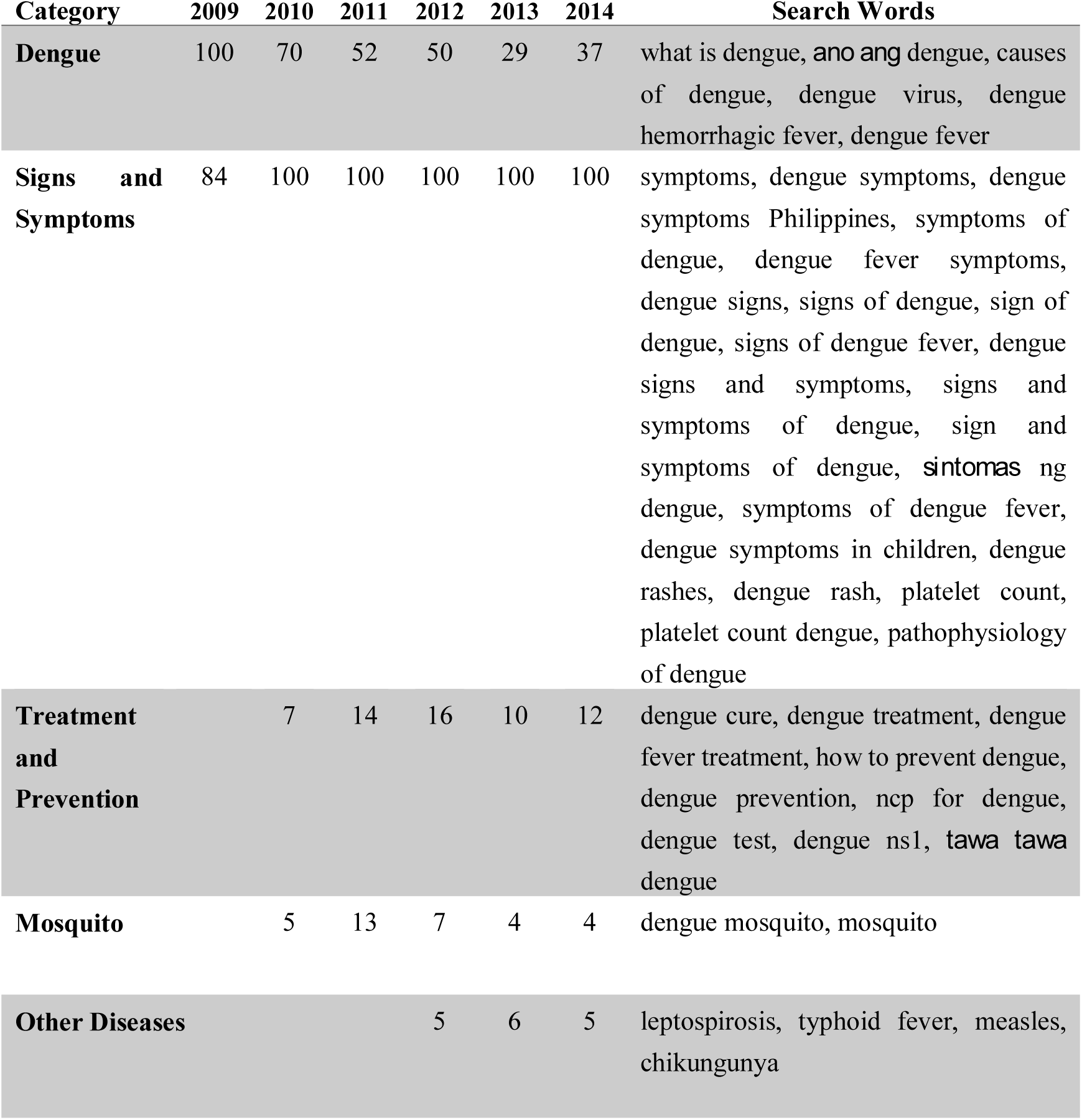
Thematic categories of top search queries related to Dengue from 2009–2014 in Metropolitan Manila. Values from 2009 – 2014 are percentages (%) based on the overall GDT values of each category per year.

Over the six-year period, search queries related to “signs and symptoms” were more visually pronounced in each word cloud (Figure 2, m-r) considering that more users searched for terms related to dengue signs and symptoms from 2010 to 2014. While some search queries were aimed at retrieving information on dengue’s signs and symptoms using search queries, such as “symptoms” and “dengue signs and symptoms”, other search queries denote specific signs and symptoms such as “dengue rash” and “platelet count.”

Some search queries were also about dengue “treatment and prevention”. A few notable search queries under this group were: (1)“ncp for dengue”, which stands for Nursing Care Plan (a reflection of the popularity of nursing as a tertiary-level course in the Philippines), (2) “dengue NS1” which concerns the dengue NS1 antigen test, and (3) “tawa-tawa dengue” which refers to Euphorbia hirta Linn, a herbal medicine believed to manage dengue.

Another category involved “mosquito” which includes search queries such as “dengue mosquito” and “mosquito”. Lastly, the “other diseases” category showed disease-specific search queries that appeared only in 2012 to 2014. “Leptospirosis” and “typhoid fever” were relevant to dengue since these occur during rainy seasons in which dengue cases tend to be more prevalent [41]. “Measles” also appeared since flooding during the rainy seasons can lead to displacement of citizens in evacuation centers where measles tends to spread rapidly [42]. “Chikungunya” was also a relevant search query due to its similarity to dengue and the considerable media attention it received in 2013 because of occasional outbreaks [42].

### 3.3. Rising and breakout search queries

A rising search query is defined by an accompanying percentage which reflects the search query’s growth in volume in a given period with respect to the preceding period [43] while breakout search query is defined by a tremendous increase in percentage. Higher percentages indicate that the search query is trending in Google [44,45]. Table 3 enumerates the list of rising and breakout search queries for each period.

**Table 3.**
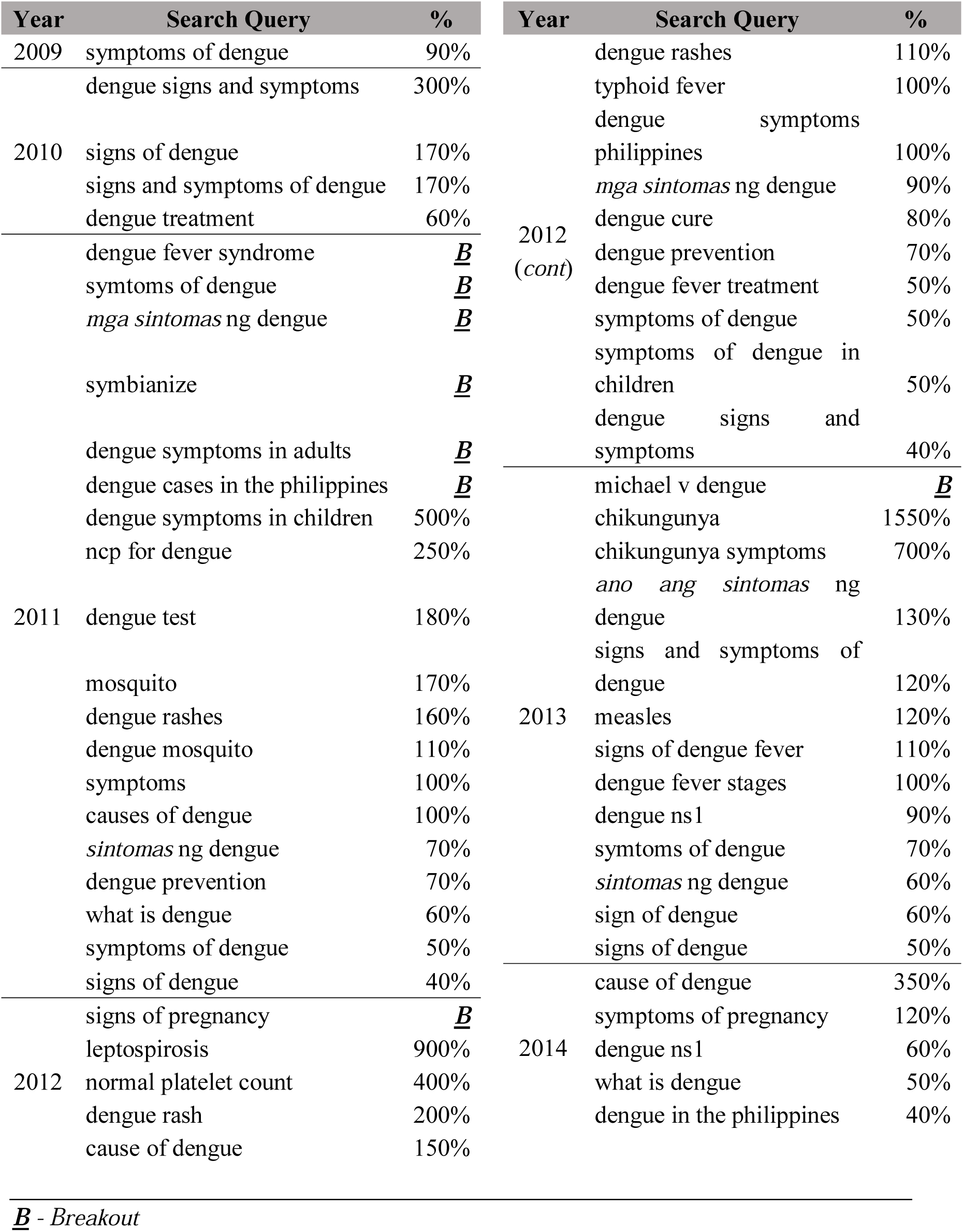
Rising and breakout search queries related to Dengue from 2009–2014 in Metropolitan Manila

Results showed that the listed rising search queries per year revolved around the five previously-mentioned thematic categories. A majority of the search queries were related to “signs and symptoms”. Although most of the search terms had a rising search percentage of 40% to 1550%, there were some that achieved breakout—search queries that grew by more than 5000% [45]. The breakout search queries from 2011 reflect finding more information about the name (e.g. ‘dengue fever syndrome’), symptoms (‘symptoms of dengue’ and ‘mga sintomas ng dengue’), and cases (‘dengue cases in the Philippines’) of dengue. Interestingly, while the search query ‘symbianize’ might be irrelevant, it achieved breakout since it is a search query related to a popular Philippine forum website where users go to search and provide health information about dengue. Another interesting breakout search query is ‘Michael V dengue’ in 2013. This situation reflects the event when Michael V, a well-known Filipino celebrity, announced on Twitter that he was diagnosed with dengue [46].

## 4. Discussion

### 4.1. Pattern of GDT and Dengue Incidence

The study was able to demonstrate the temporal relatedness of GDT to dengue by adapting the adjustment procedure [30,31]. Previous studies that assessed GDT and dengue did not consider adjusting their GDT values because it used monthly rather than weekly observed values. However, one study processed GDT data by multiplying the values to a certain factor in order to achieve almost similar unit scales to the reported dengue cases [28]. Although this is understandable, most dengue endemic Asian countries such as the Philippines follow the weekly reporting of total dengue cases [48]. Therefore, integrating a weekly temporal scale would be more appropriate in predicting of future dengue outbreaks. The use of a weekly temporal scale is of utmost significance for implementing necessary health interventions or measures. It is worth noting that GDT generates relative values and is limited to a given time range, where it identifies the highest query volume to generate weekly GDT values. This is why GDT values assigned to a specific week generates different values when compared using two different time periods (e.g. one year versus two years). Therefore, appropriately adjusting GDT values is highly recommended. In the study, it clearly demonstrated the importance of adjusting when the temporal pattern and correlation coefficient were compared from non-adjusted GDT values. It was shown that the adjusted GDT captured the correct temporal trend, resulting to a higher association. Furthermore if dengue incidence was adjusted, it showed that it capture the non-adjusted GDT trend and higher correlation coefficients. This strengthens the previous recommendation that researchers conducting identical studies in the future must consider adjusting GDT values appropriately in order to correctly estimate the correct association of reported cases or incidence.

The cross-correlation results revealed that GDT has a delayed effect (Lag 1) to the reported dengue incidence. This delayed effect may stem towards the user’s behavior of self-diagnoses of one’s physical condition before going to the hospital for possible admission and management of the disease. This hypothesis corroborates with our thematic analysis where majority of search queries of dengue refer to its ‘signs and symptoms’. It was claimed that an increase of health seeking behavior from internet users would pertain the surrounding environment being affected by the disease [28]. Due to the untimely release of traditional disease surveillance reports to the public, such cannot be used for quasi-real time disease monitoring. Therefore, GDT could act as a potential compliment for real-time disease monitoring tools in order to alert public health officials for early phases of an outbreak [48]. However, this approach has its limitations or drawbacks as exemplified when Google Trends was assessed during the influenza pandemic of 2009, the 2012/2013 flu epidemic season in the US and, the Ebola pandemic in Africa [48,49]. These studies outlined that Google Trends platform is limited to the proportion of the population who use the internet to obtain health-related information [48], intense media coverage [49-51] and algorithm dynamics [50-52]. Moreover, the application of GDT may only be suited in areas with high dengue incidence [29]. Thus, employing GDT in Metropolitan Manila can be well-suited due to its high occurrence of the disease and huge internet accessibility of users. Furthermore, the inclusion of meteorological variables and novel modeling approaches may further strengthen the prediction of dengue incidence in this highly-urbanized region in the Philippines [53]. Since the study was only limited in investigating in one region in the Philippines, our interpretation cannot be the same in other regions with a large proportion of users situated in rural areas presumably with no internet access. The selective application of GDT may only be a powerful tool to urban diseases such as dengue where there is available internet use.

### 4.2. Search Query Behavior towards Dengue

The results showed that most of the search query activity for dengue came from the cities of Makati, Quezon, and Manila. These cities also consistently showed high GDT values across all time periods. These three cities share common characteristics that resulted for such outcome. One of the characteristics could be the cities’ high population density that contributes to the presence of a high proportion of internet users. Previous works in the US suggest that high population density correlates with more internet users [51,54]. Another characteristic would the presence of major business districts, hospitals, and educational institutions in these cities. The presence of these establishments indicates urbanization. A study from 28 Asian countries, including the Philippines, showed that urbanization is related to higher internet penetration [55]. As of January 2018, the internet penetration rate in the Philippines is 63% which translates to 67 million users [56], majority of which are residents of urban areas such as Metropolitan Manila. In general, people living in urban areas with high population density tend to have more internet access and this facilitates greater health information seeking on the internet.

Based on the thematic analysis of the search queries, it can be inferred that users were mostly performing health information seeking on dengue. Specifically, “top”, “rising”, and “breakout” search queries mostly consist of search queries that reflect seeking information about dengue including its signs and symptoms. This is somewhat expected considering that the internet acts an enabler for people to search for health information [57]. At the same time, the findings reflect the knowledge gap hypothesis [58] wherein that people from an urban area, like Metropolitan Manila (regardless of socioeconomic background), can reduce their knowledge gaps about health-related queries (dengue) through the internet.

While the findings indicate that the internet acts as an empowering technology for people to gain more knowledge about dengue, a consequence of health information seeking over the internet is self-diagnoses of dengue without proper medical consultation [59,60]. Such situation might lead to “cyberchondriasis” where people start to develop health anxiety [61]. Scholars have advised that searching for disease-specific search queries may not correspond to true illness [15]. Considering that some of dengue’s signs and symptoms might overlap with other diseases, such as chikungunya and measles, the values reflected in Google Trends might not fully indicate disease-specific occurrence and might just reflect greater health anxiety among people.

Another way of interpreting the findings is through the lens of the agenda setting theory [62]. The theory indicates that the media (news outlets) has the influence on placing importance to topics of public agenda. This could explain some of the breakout search queries since it might be a result of high media coverage of an incident that reflect the search query. For instance, media reports of increasing dengue cases in Metropolitan Manila and other regions in 2011 [63] might have led people to perform more search queries related to dengue signs and symptoms than in 2010. Moreover, the proliferation of news related to the dengue diagnosis of Michael V, a Filipino celebrity [46], in 2013 might have spiked interests to search for “Michael V dengue” on Google during that year.

## 5. Conclusions

The study was able to demonstrate the high temporal relationship between the weekly patterns of GDT and dengue. It was able to reveal the necessary adjusting procedures in order to capture the correct temporal trend and, likewise, produce higher correlation coefficients between the two values. With this, the study was able to demonstrate delayed (lag) effect of GDT towards dengue incidence, therefore, has the potential to be utilized in detecting future disease outbreaks and patterns. Although Google Trends is limited to the proportion of internet users and may possibly be only suitable in areas with high disease incidence and internet penetration, it undoubtedly has the capacity to assist the traditional disease surveillance. And while it is unclear how these limitations can be addressed, it can be used as a core component for future studies regarding the utilization of Google Trends for disease surveillance.

The study also revealed health-seeking behaviors of the population by evaluating dengue-related search queries in Metropolitan Manila from 2009-2014. Thematic analyses revealed 5 categories namely; (a) dengue, (b) sign and symptoms of dengue, (c) treatment and prevention, (d) mosquito and (e) other diseases. Further analysis showed that majority of the search activity pertains to ‘signs and symptoms’ of dengue. These findings indicate how the internet acts as an empowering technology in gaining knowledge about dengue. However, users might also utilize this technology in conducting self-diagnoses without proper medical consultation. In addition to this, the high dengue search activity of the population may be attributed to the influence of media which could have led the population to perform more search queries related to dengue’s signs and symptoms.

## Acknowledgments

This paper is dedicated to the formation of the Nursing Science Research Department of St. Luke’s College of Nursing – Trinity University 10 years ago. The three authors (HTH, TMC and JRD) were considered as key individuals for its establishment. This work is funded by the JSPS Grant-in-Aid for Scientific Research (16H05750, 17H01624, 17K18906), JSPS Bilateral Joint Research Projects, and Leading Academia in Marine and Environmental Pollution Research – Ehime University (Y29-1-8; 30-04)

## Author Contributions

Concept and Design: HTH TMC JRR and KW. Data Collection and Pre-processing: TMC, KMV, LFTH, JDRC, JRR and HTH. Data Analysis and Interpretation: TMC, KMV, LFTH, JDRC, JRR and HTH. Drafting of the manuscript: HTH, TMC, JDRC, JRR and KW. All authors read and approved the final manuscript.

## Supplemental Material

**Figure S1.** Administrative Boundaries of Metropolitan Manila Cities

**Table S1.** Cross-correlation analysis (R^2^) of Dengue incidence and Google Dengue Trends.

